# Quantum kernel support vector machines for trabecular bone classification: comparing feature reduction strategies on synthetic micro-CT data

**DOI:** 10.64898/2026.05.04.722627

**Authors:** Isabella Florez, Ahmed Farhat, James Le Houx, Edoardo Altamura, Gianluca Tozzi

## Abstract

Quantum kernel methods offer a potential advantage for classification tasks in high-dimensional feature spaces, yet their practical benefit critically depends on how input features are prepared. We compare five dimensionality reduction strategies—principal component analysis (PCA), Gaussian random projection (RP Gaussian), sparse random projection (RP Sparse), partial least squares (PLS), and uniform manifold approximation and projection (UMAP) — as pre-processing steps for quantum kernel support vector machines (SVMs) applied to trabecular bone classification from synthetic micro-computed tomography (micro-CT) data. Using a custom procedural generator based on Gaussian random field zero-crossings, we produced 500 synthetic trabecular bone volumes with controlled morphometric properties such as bone volume fraction (BV/TV), trabecular thickness (Tb.Th), number (Tb.N) and spacing (Tb.Sp). Texture features extracted from grayscale slices are reduced to 8-dimensional quantum circuit inputs via each method, then classified using both classical radial basis function (RBF)-SVMs and quantum kernel SVMs with ZZ feature maps on a statevector simulator, both evaluated with 5 × 5 repeated stratified cross-validation (25 folds). Our results show that UMAP is the only reduction method where the quantum kernel remains competitive with the classical baseline. Under repeated cross-validation, UMAP showed a +0.032 accuracy gap favouring the quantum kernel (Dietterich 5 × 2 CV *p* = 0.177); however, validation on 10 fully independent datasets—each with independently generated samples, separate reduction fits, and separate kernel matrices — reversed the sign to −0.030 (paired *t*-test *p* = 0.123; Wilcoxon *p* = 0.193; quantum wins 3/10 datasets), indicating that the apparent advantage was likely inflated by fold dependence. Nevertheless, UMAP’s gap remains small and non-significant in both analyses, whereas all linear methods (PCA, RP Gaussian, PLS) show substantial quantum deficits of −0.090 to −0.116 across BV/TV classification, with PCA and PLS remaining significant under corrected tests (5 × 2 CV *p* = 0.004 and *p* = 0.007 respectively). We additionally evaluate quantum kernel ridge regression for continuous morphometric prediction, finding that ZZ quantum kernels fail uniformly at regression (negative *R*^2^ for all methods except PLS at 4 qubits), suggesting that the ZZ kernel captures decision boundaries but not smooth metric structure. These findings provide practical guidance for feature engineering in near-term quantum machine learning pipelines and demonstrate that the choice of dimensionality reduction can determine whether quantum kernels remain competitive with classical baselines.

## 1 Introduction

Quantum machine learning (QML) has attracted significant interest as a potential application of near-term quantum computers [1]. Among QML approaches, quantum kernel methods are particularly appealing because they leverage quantum circuits to compute kernel functions in exponentially large Hilbert spaces while relying on well-understood classical support vector machine (SVM) optimisation for the learning step [2]. However, the practical benefit of quantum kernels over classical alternatives remains an open question, with recent work suggesting that advantages depend sensitively on data properties and feature preparation [3]. A fundamental obstacle is exponential concentration: Thanasilp et al. [4] proved that quantum kernel values can concentrate exponentially (in the number of qubits) towards a fixed value under conditions including high embedding expressivity, entanglement, global measurements, and noise, rendering the Gram matrix uninformative with polynomial measurement budgets.

Havlicek et al. [2] introduced quantum kernel estimation using parameterised circuits, demonstrating that quantum feature maps can define kernel functions which are unmanageable for classical computation. Schuld and Killoran [5] formalised the connection between quantum feature maps and reproducing kernel Hilbert spaces, establishing a theoretical framework for quantum kernel design. Huang et al. [3] subsequently showed that the advantage of quantum kernels depends on the geometric properties of the data: they can outperform classical methods only when the data exhibit structural alignment with the quantum feature space. These foundational results motivate careful attention to how data is prepared before encoding into quantum circuits.

More recently, Jiang and Otten [6] demonstrated that encoding strategy dominates quantum kernel performance, with over 30% accuracy variation across encoding choices on eight real-world datasets. Their variational kernel framework with kernel-target alignment optimisation achieved consistent improvements but notably used adaptive parameterisation that is considered a more flexible setting than our fixed ZZ feature map, which makes UMAP’s competitiveness under a constrained encoding particularly notable. Nooblath et al. [7] applied PCA-reduced features to quantum kernels with RealAmplitudes and EfficientSU2 approach, finding that full entanglement outperformed linear and circular topologies (80.4% vs 76.0% for RBF baseline), and that entanglement strategy interacts with dimensionality.

The mismatch between classical feature dimensionality and available qubit counts makes dimensionality reduction a practical necessity for near-term QML. Shinde and Nurminen [8] systematically evaluated the influence of reduction methods such as PCA, singular value decomposition (SVD), autoencoders and t-distributed stochastic neighbour embedding (t-SNE) on quantum model performance, reporting 14–48% accuracy differences depending on method choice. They found that quantum neural networks generally benefit from reduction, but QSVM performance often degrades. Critically, their study did not evaluate UMAP or random projections, leaving open whether nonlinear manifold methods behave differently. Slabbert and Petruccione [9] used a ResNet10 autoencoder to reduce image features to 64 dimensions for amplitude-encoded 6-qubit QSVMs, identifying the classical preprocessing stage as the performance bottleneck. However, they evaluated only a single extraction method without comparing alternatives.

Recent theoretical work has clarified UMAP’s algorithmic nature. Yang [10] proved that UMAP performs spectral clustering on the fuzzy *k*-nearest neighbour graph, establishing a chain of equivalences: UMAP’s stochastic optimisation with negative sampling is a contrastive learning objective, which is equivalent to spectral clustering via the graph Laplacian. This result shows that UMAP embeddings are determined by the spectral properties of the neighbourhood graph rather than by global variance or random projections; a distinction that may have direct consequences for quantum kernel compatibility.

Trabecular bone microarchitecture, assessed via micro-computed tomography (micro-CT), is a critical determinant of bone quality and fracture risk [11]. Standard morphometric parameters such as bone volume fraction (BV/TV), trabecular thickness (Tb.Th), trabecular number (Tb.N), and trabecular separation (Tb.Sp) are widely employed to characterise this architecture but require segmented 3D volumes. Bouxsein et al. [11] established guidelines for micro-CT assessment of trabecular bone, defining these standard morphometric parameters. Texture analysis of bone images has a long history in medical imaging, with grey-level co-occurrence matrices (GLCM) [12] providing established second-order spatial features. Automated classification of bone quality from image texture features could enable faster clinical assessment, making this an attractive test case for machine learning methods including quantum approaches.

A key challenge for quantum kernel methods is dimensionality reduction: quantum circuits with *n* qubits accept *n*-dimensional inputs, typically *n* ≤ 12 on current simulators, yet image-derived feature vectors may contain hundreds of dimensions. The choice of reduction method determines what information reaches the quantum circuit and therefore whether the quantum kernel can outperform classical alternatives. Recent work has examined classical-quantum preprocessing pipelines for image classification [9], and the influence of reduction methods on quantum model performance [8], but no systematic comparison has evaluated multiple reduction families — including linear, random, supervised, and nonlinear — against each other for quantum kernel SVMs.

In this work, we introduce a novel procedural generator for synthetic trabecular bone micro-CT based on zero-crossings of Gaussian random fields, capable of producing topology spanning the spectrum from rod-like to plate-like architecture with controllable morphometric parameters. This is clinically relevant because plate-like trabecular bone, consisting of flat, sheet-like structures, is associated with higher mechanical strength, while rod-like trabecular bone, consisting of cylindrical, column-like structures, is associated with lower bone strength and the conversion of plates into rods is a hallmark of age-related bone loss [13]. A systematic comparison of five dimensionality reduction methods — PCA, RP Gaussian, RP Sparse, PLS, and UMAP — as quantum kernel input preparation was evaluated on both classification and regression tasks. Our results show that UMAP is the only reduction method where the quantum kernel remains competitive with the classical baseline for BV/TV classification, while all linear methods (PCA, RP Gaussian, PLS) show substantial quantum deficits. This in turn provides clear and actionable guidance for quantum feature engineering.

## 2 Methods

### 2.1 Synthetic Trabecular Bone Generation

We developed a procedural volumetric generator that produces synthetic micro-CT images of trabecular bone. The generation process begins by constructing a 3D isotropic Gaussian random field [14]: a white noise volume is smoothed with a Gaussian kernel (*σ* = 3.0 voxels) to produce a spatially correlated scalar field with cellular-like topology. This field is then deformed by smooth elastic displacement fields (amplitude = 1.5, correlation length = 10.0 voxels) to introduce the natural irregularity characteristic of biological tissue.

Bone structure is extracted from the deformed field via zero-crossing wall detection: voxels where |field| *< τ* are labelled as bone, where *τ* is a wall thickness parameter. The zero level-set of a smooth Gaussian random field forms a connected surface network that is topologically equivalent to trabecular plates, making this approach naturally suited to plate-dominant bone architecture. By varying the plate/rod weighting parameter, the generator can produce topology spanning the continuum from rod-dominant to plate-dominant structures, although the present study targets a plate/rod weight of 0.7/0.3 to reflect the plate-dominant architecture typical of healthy vertebral trabecular bone. The parameter *τ* is calibrated via binary search to achieve a target BV/TV.

The resulting binary bone masks are then converted to realistic synthetic micro-CT grayscale through distance-transform based intensity fill [15] (bone mean = 95, marrow mean = 20), partial volume blurring (*σ* = 1.8), additive Gaussian noise (*σ* = 3.0), and background texture noise (*σ* = 1.0). Morphometric targets (BV/TV, Tb.Th, Tb.Sp) were sampled from a multivariate Gaussian fitted to 18 volumes of interest (VOIs) extracted from a previously published micro-CT dataset of porcine vertebrae scanned at 39 µm isotropic resolution [16, 17], preserving inter-parameter correlations. Quality gates rejected samples with *>* 8% BV/TV relative error. The *z*-dimension (40 slices) was chosen to maintain computational tractability for quantum kernel matrix computation while preserving sufficient volumetric context for meaningful texture extraction. Key generator parameters are listed in Table 1.

**Table 1:** Generator parameters.

| Parameter | Value |
| --- | --- |
| Version | v15.3 zero-crossing |
| Base sigma | 3.0 voxels |
| Target BV/TV range | [0.10, 0.40] |
| Target Tb.Th | 180 $\mu\text{m}$ |
| Anisotropy ratio | 1.0 (isotropic) |
| Warp amplitude / sigma | 1.5 / 10.0 voxels |
| Plate / rod weight | 0.7 / 0.3 |
| Bone / marrow mean | 95 / 20 |
| Blur sigma | 1.8 |
| Noise $\sigma$ | 3.0 |
| Voxel size | 39 $\mu\text{m}$ isotropic |
| Volume size | $128 \times 128 \times 40$ voxels |

### 2.2 Feature Extraction

From each synthetic volume, we extracted texture features from the mid-slice grayscale image resized to 64 × 64 pixels. While full volumetric feature extraction would exploit the 3D microarchitecture directly, we deliberately restrict to 2D slice-based features for two reasons: first, it mirrors the clinical scenario where a single representative slice may be available or preferred for rapid screening; second, it provides a more challenging test case for the classifiers — if quantum kernels can demonstrate an advantage on limited 2D texture information, the finding is likely to hold with richer 3D features.

The feature set comprises 31 first-order statistical descriptors capturing the intensity distribution of each slice, including central tendency measures (mean, median), dispersion measures (standard deviation, min, max, 25th/75th percentiles), shape measures (skewness, kurtosis), a 16-bin normalised histogram, and gradient magnitude statistics (mean, standard deviation, P75, P95) along with local variance statistics. These are complemented by 12 second-order texture descriptors derived from grey-level co-occurrence matrices (GLCM) [12] computed at 32 quantisation levels: contrast, dissimilarity, homogeneity, energy, correlation, and angular second moment, each summarised as the mean and standard deviation over 2 distances × 4 angles. The combined 43-dimensional feature vector thus captures both the global intensity profile and the local spatial texture patterns relevant to trabecular microarchitecture.

### 2.3 Dimensionality Reduction

Since quantum circuits with *n* qubits accept *n*-dimensional inputs, the 43-dimensional feature vectors must be reduced before encoding. We compared five reduction methods spanning four methodological families to understand how each interacts with quantum kernel computation. PCA provides a linear, variance-maximising baseline by projecting onto principal components. Two random projection methods were included: RP Gaussian, which uses a dense Gaussian random matrix and approximately preserves pairwise distances via the Johnson–Lindenstrauss lemma [18], and RP Sparse, which uses a sparse random matrix [19] (density 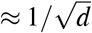) for computational efficiency. PLS [20] takes a supervised approach, maximising the covariance between features and morphometric targets. Finally, UMAP [21] provides a nonlinear alternative that preserves local neighbourhood structure through manifold learning.

All methods reduced the components from 43 to 16 (except PLS: 4 components, limited by the number of target variables). Features were standardised using RobustScaler before reduction. For quantum kernel input, reduced features were rescaled to [0, *π*] via min-max scaling to match the ZZ feature map’s periodic encoding. For classical SVM input, features were rescaled to [0, 1].

### 2.4 Classical and Quantum SVM

With the reduced feature representations in hand, we evaluated binary classification on two tasks: BV/TV (sparse vs dense bone, median split at BV/TV = 0.330) and Tb.N (low vs high trabecular number, median split at Tb.N = 2.90/mm). For the classical baseline, we trained an RBF-kernel SVM (*C* = 1.0, *γ* = scale) and evaluated it via 5 × 5 repeated stratified cross-validation, yielding 25 accuracy estimates per method across all 500 samples. For the quantum kernel, we constructed a ZZFeatureMap [2] with 2 repetitions and linear entanglement on 8 qubits (4 qubits for PLS due to its fewer components) and computed the quantum kernel via Qiskit’s FidelityQuantumKernel using statevector simulation [22]. To ensure a fair comparison, the full 500 × 500 quantum kernel matrix was precomputed once per reduction method, requiring approximately 140 minutes per matrix. The same 5 × 5 repeated stratified cross-validation was then performed by indexing into this precomputed matrix, so that both classical and quantum classifiers were evaluated on identical folds with matched *±* std estimates.

Both classifiers were evaluated on identical fold partitions using RepeatedStratifiedKFold with a fixed random seed (seed = 42), ensuring valid pairing for statistical comparison. We note that fold estimates in repeated cross-validation are not fully independent, as training sets overlap both within and across repeats. To account for this dependence, we report three statistical tests: (i) the uncorrected Wilcoxon signed-rank test [23] on 25 paired fold accuracies, (ii) the Nadeau–Bengio corrected paired *t*-test [24], which inflates variance by a factor of (1 + *n*_test_*/n*_train_) to account for training set overlap, and (iii) the Dietterich 5 × 2 CV paired *t*-test [25], which uses five independent 50/50 splits with df = 5. Alternative resampling strategies such as bootstrapping could also be used to obtain more robust uncertainty estimates; the independent dataset validation described below provides a complementary approach that avoids fold-dependence entirely.

### 2.5 Independent Dataset Validation

Repeated cross-validation, even with corrected tests, evaluates a single dataset and therefore cannot fully disentangle classifier differences from dataset-specific artefacts. To address this, we performed an independent dataset validation for UMAP-based BV/TV classification — the only reduction method that showed quantum competitiveness in the repeated CV analysis. Ten fully independent datasets of 500 samples each were generated using the same procedural bone generator with independent random seeds. For each dataset, the entire pipeline was executed independently: feature extraction, UMAP fitting, feature scaling, quantum kernel matrix computation, and SVM evaluation via a single stratified 80/20 train-test split. This design eliminates all sources of fold dependence — each accuracy estimate derives from a completely separate dataset with no shared training samples, no shared reduction fit, and no shared kernel matrix. Total computation time for the 10 independent kernel matrices was approximately 27.7 hours. A paired *t*-test (df = 9) and Wilcoxon signed-rank test were applied to the 10 paired accuracy differences, with significance assessed at *α* = 0.05.

### 2.6 Quantum Kernel Ridge Regression

Beyond classification, we sought to determine whether quantum kernels capture smooth metric structure suitable for continuous prediction. Using the same precomputed ZZ kernel matrices, we performed quantum kernel ridge regression (KRR) [26] for continuous BV/TV prediction via scikit-learn’s Kernel-Ridge [27]. Regularisation *α* was selected by grid search over [0.001, 0.01, 0.1, 1.0, 10.0] on a stratified 80/20 split (400 train, 100 test). Classical Ridge regression with leave-one-out cross-validation (RidgeCV) served as baseline, allowing us to assess whether the quantum kernel’s utility extends from decision boundary construction to smooth interpolation.

## 3 Results

### 3.1 Dataset Characteristics

Tb.N was derived as 1*/*(Tb.Th + Tb.Sp). All values fall within the quality gates specified in Section 2.1. All 500 samples achieved LCC_raw_ = 1.000 (perfect connectivity), confirming the zero-crossing generator produces fully connected plate networks. Dataset morphometric distribution is reported in Table 2.

**Table 2:**
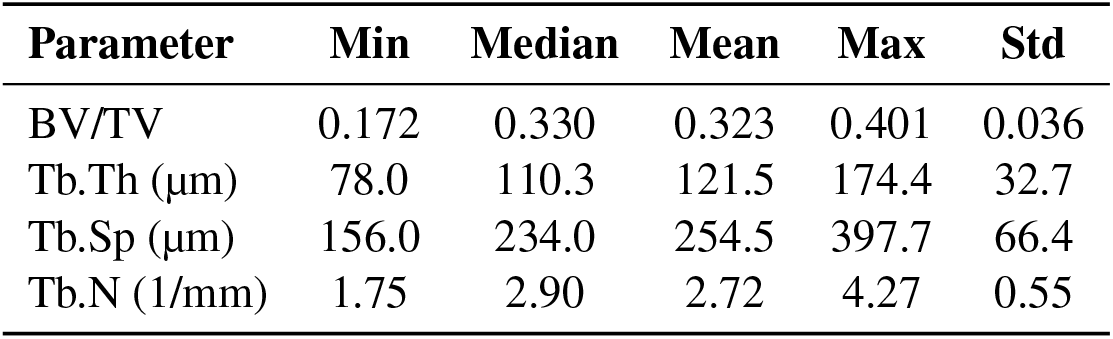
Dataset morphometric distributions (*n* = 500).

### 3.2 Reduction Quality Validation

To validate that the reduced representations retained meaningful morphometric information, we measured downstream Ridge regression *R*^2^ predicting all four morphometric targets from the reduced features (Table 3). All linear methods achieved comparable *R*^2^ *≈* 0.91, confirming that the 43 texture features contain substantial morphometric information that survived linear compression. UMAP scored lower (0.872), consistent with its nonlinear structure; Ridge regression cannot fully exploit UMAP’s manifold-preserving geometry, which is better suited to nonlinear downstream models such as quantum kernels.

**Table 3:** Downstream Ridge regression *R*^2^ by reduction method.

| Method | Mean | BV/TV | Tb.Th | Tb.N | Tb.Sp |
| --- | --- | --- | --- | --- | --- |
| RP Sparse | 0.911 | 0.849 | 0.922 | 0.939 | 0.936 |
| PCA | 0.910 | 0.836 | 0.923 | 0.944 | 0.936 |
| RP Gauss. | 0.909 | 0.842 | 0.928 | 0.932 | 0.934 |
| PLS | 0.909 | 0.832 | 0.925 | 0.942 | 0.936 |
| UMAP | 0.872 | 0.818 | 0.901 | 0.868 | 0.899 |

### 3.3 BV/TV Classification

Table 4 presents BV/TV classification accuracy with both classical and quantum SVMs evaluated under identical 5 × 5 repeated stratified cross-validation. RP Sparse was omitted due to excessive runtime (8 hours) and poor single-split performance in preliminary experiments. Rows are sorted by quantum– classical gap. UMAP is the only method where the quantum kernel exceeds the classical baseline (+0.032), with the quantum kernel outperforming the classical baseline on 19 of 25 folds. However, since fold estimates in repeated cross-validation are not fully independent — training sets overlap both within and across repeats — we additionally performed two corrected statistical tests. The Nadeau–Bengio corrected paired *t*-test, which inflates variance to account for fold dependence, yields *p* = 0.168. The Dietterich 5 × 2 CV paired *t*-test, which uses five independent 50/50 splits with proper degrees of freedom (df = 5), yields *p* = 0.177 with the quantum kernel winning 9 of 10 folds. We therefore report the UMAP quantum–classical gap under repeated CV as a consistent positive trend that does not reach statistical significance under corrected tests.

**Table 4:**
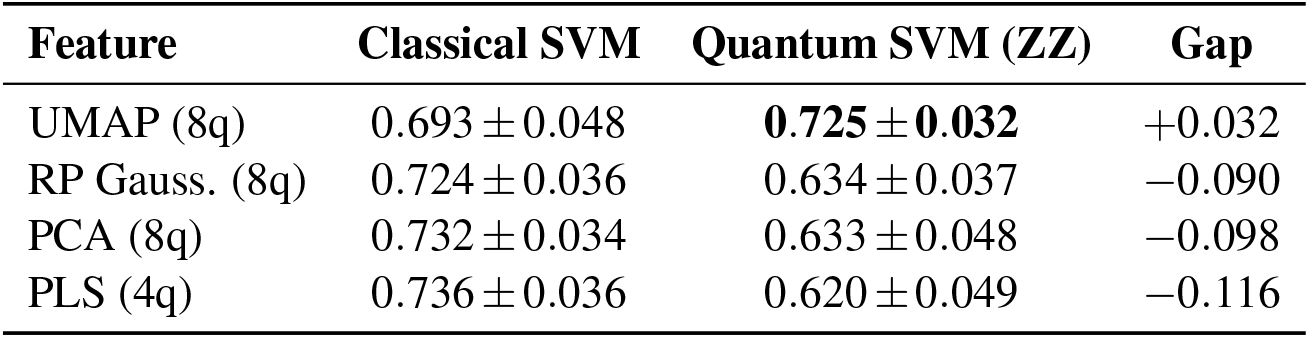
BV/TV classification — 5 × 5 repeated stratified CV.

| Feature | Classical SVM | Quantum SVM (ZZ) | Gap |
| --- | --- | --- | --- |
| UMAP (8q) | $0.693 \pm 0.048$ | **$0.725 \pm 0.032$** | +0.032 |
| RP Gauss. (8q) | $0.724 \pm 0.036$ | $0.634 \pm 0.037$ | −0.090 |
| PCA (8q) | $0.732 \pm 0.034$ | $0.633 \pm 0.048$ | −0.098 |
| PLS (4q) | $0.736 \pm 0.036$ | $0.620 \pm 0.049$ | −0.116 |

The three linear methods (RP Gaussian, PCA, PLS) all show substantial quantum deficits (− 0.090 to − 0.116). Under the Dietterich 5 × 2 CV test, the PCA deficit remains significant (*p* = 0.004) and PLS remains significant (*p* = 0.007), while RP Gaussian drops to non-significant (*p* = 0.123), likely due to the test’s limited degrees of freedom (df = 5). Under the Nadeau–Bengio corrected test, all three linear methods remain highly significant (all *p <* 0.001). PLS performs worst despite its supervised feature construction. Note that PLS uses 4 qubits rather than 8 due to its component limit, making direct comparison with the 8-qubit methods approximate.

### 3.4 Independent Dataset Validation

Table 5 presents the results of the independent dataset validation for UMAP-based BV/TV classification. Across 10 fully independent datasets of 500 samples each, the classical SVM achieved a mean accuracy of 0.721 *±* 0.045 and the quantum SVM achieved 0.691 *±* 0.048, yielding a mean gap of −0.030 (quantum deficit). The quantum kernel outperformed the classical baseline on only 3 of 10 datasets. Neither the paired *t*-test (*t* = −1.70, df = 9, *p* = 0.123) nor the Wilcoxon signed-rank test (*p* = 0.193) reached significance. Figure 3 visualises the paired results, showing that while individual datasets occasionally favour the quantum kernel, the majority of lines slope downward from classical to quantum.

**Table 5:** UMAP BV/TV classification — independent dataset validation (*k* = 10).

| Dataset | Classical | Quantum | Gap | Q wins |
| --- | --- | --- | --- | --- |
| 1 | 0.700 | 0.740 | +0.040 | ✓ |
| 2 | 0.710 | 0.700 | −0.010 |  |
| 3 | 0.760 | 0.780 | +0.020 | ✓ |
| 4 | 0.620 | 0.660 | +0.040 | ✓ |
| 5 | 0.730 | 0.620 | −0.110 |  |
| 6 | 0.710 | 0.700 | −0.010 |  |
| 7 | 0.770 | 0.660 | −0.110 |  |
| 8 | 0.710 | 0.670 | −0.040 |  |
| 9 | 0.780 | 0.730 | −0.050 |  |
| 10 | 0.720 | 0.650 | −0.070 |  |
| **Mean** | **0.721 ± 0.045** | **0.691 ± 0.048** | **−0.030** | **3/10** |

**Figure 1.**
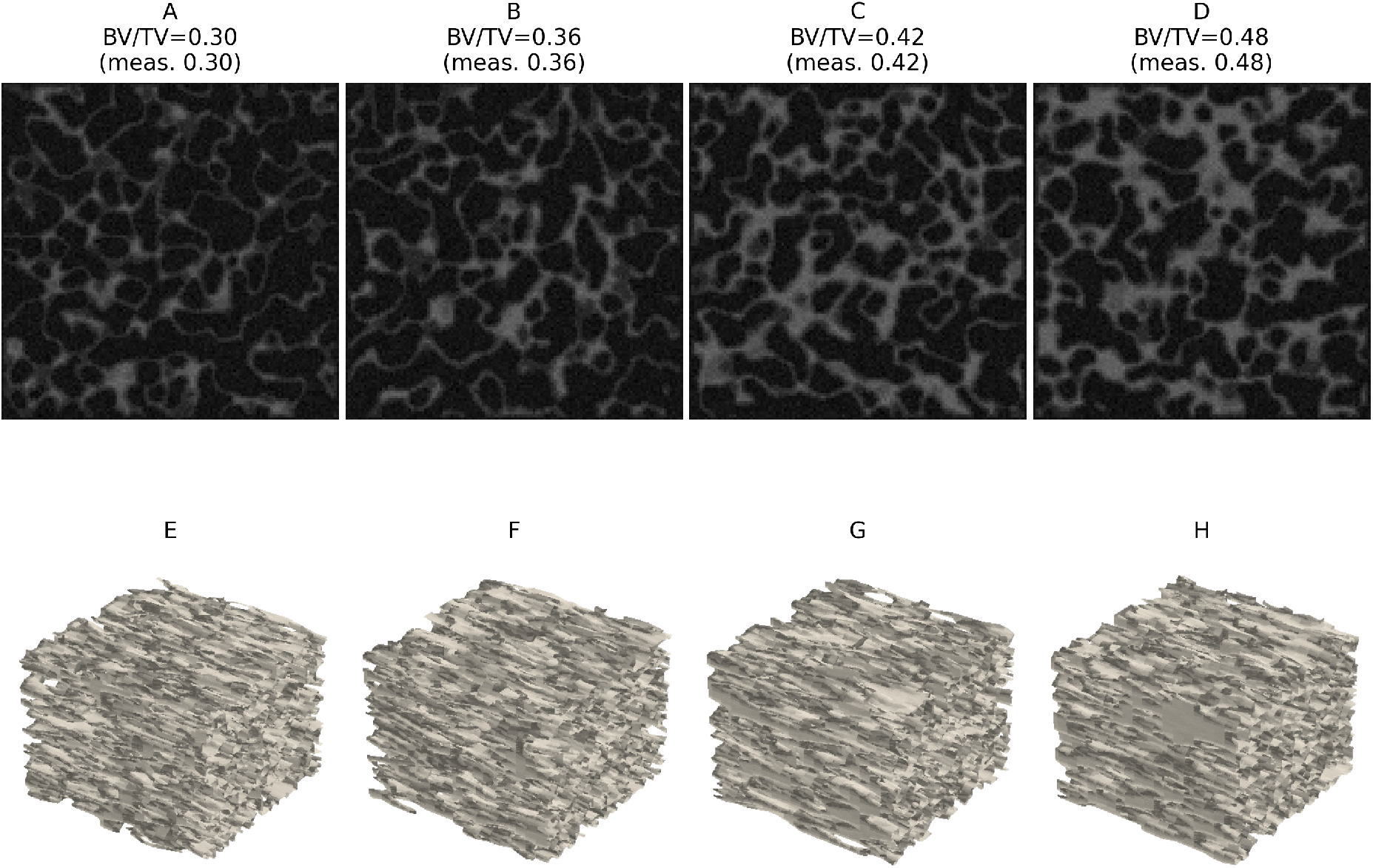
Synthetic trabecular bone generated using the zero-crossing Gaussian random field model with increasing target BV/TV. (A–D) Grayscale mid-slice cross-sections at BV/TV = 0.30, 0.36, 0.42, and 0.48 respectively. (E–H) Three-dimensional isosurface renderings of the corresponding synthetic volumes, illustrating the transition from rod-dominant to plate-dominant trabecular architecture. All samples are shown at 39 µm isotropic voxel resolution with a plate/rod weight of 0.7/0.3.

**Figure 2.**
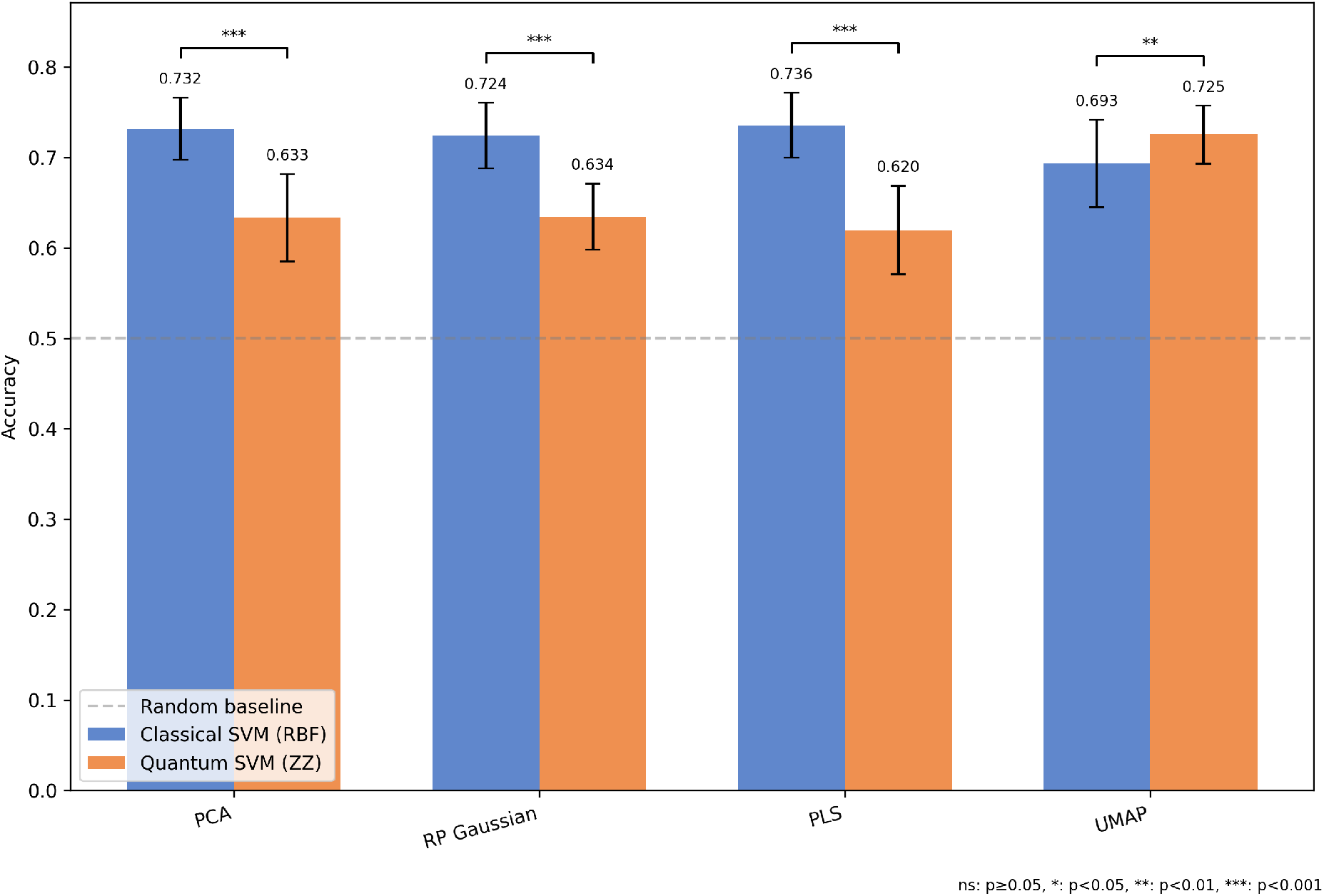
BV/TV classification accuracy for classical and quantum SVMs across dimensionality reduction methods. Blue bars represent classical RBF-SVM results, whereas orange bars represent quantum SVM results with a ZZ feature map. Error bars show *±*1 standard deviation across 25 folds obtained from 5 × 5 repeated stratified cross-validation. The dashed line indicates the random baseline (0.50). Brackets show significance from the Dietterich 5 × 2 CV paired *t*-test (df = 5). The table below reports *p*-values from three statistical tests: uncorrected Wilcoxon signed-rank, Nadeau–Bengio corrected paired *t*-test, and Dietterich 5 × 2 CV test. UMAP is the only method where the quantum kernel matches or exceeds the classical baseline under repeated CV, though this finding is not confirmed by independent dataset validation (see Section 3.4).

**Figure 3.**
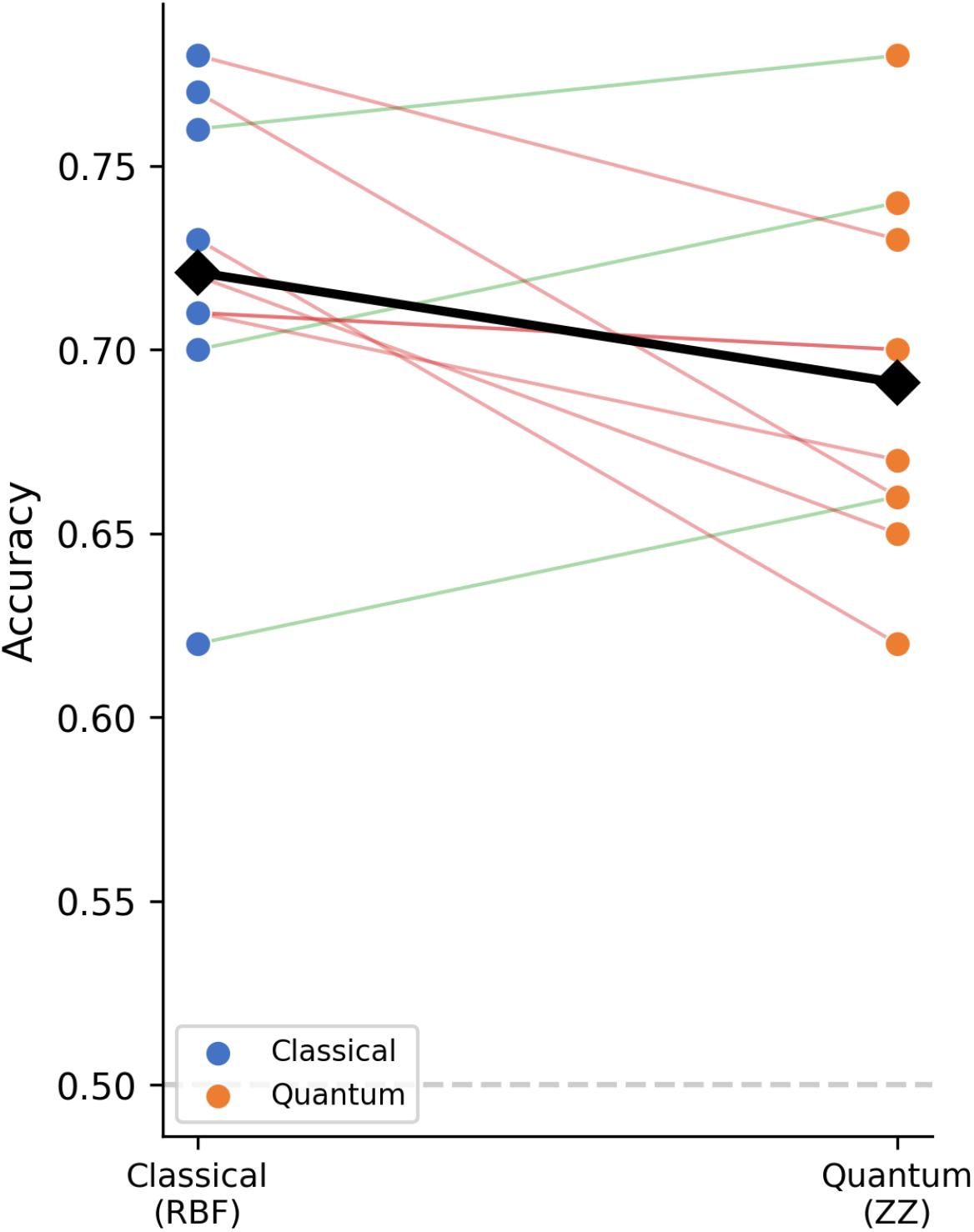
Paired comparison of classical (RBF) and quantum (ZZ) SVM accuracy for UMAP-based BV/TV classification across 10 fully independent datasets. Each thin line connects a paired classical– quantum result for one dataset; green lines indicate quantum wins (3/10) and pink lines indicate classical wins (7/10). The bold black line connects the group means (0.721 → 0.691, gap =− 0.030). The dashed grey line marks the random baseline (0.50). The mean gap is not statistically significant (paired *t*-test *p* = 0.123; Wilcoxon *p* = 0.193).

**Figure 4.**
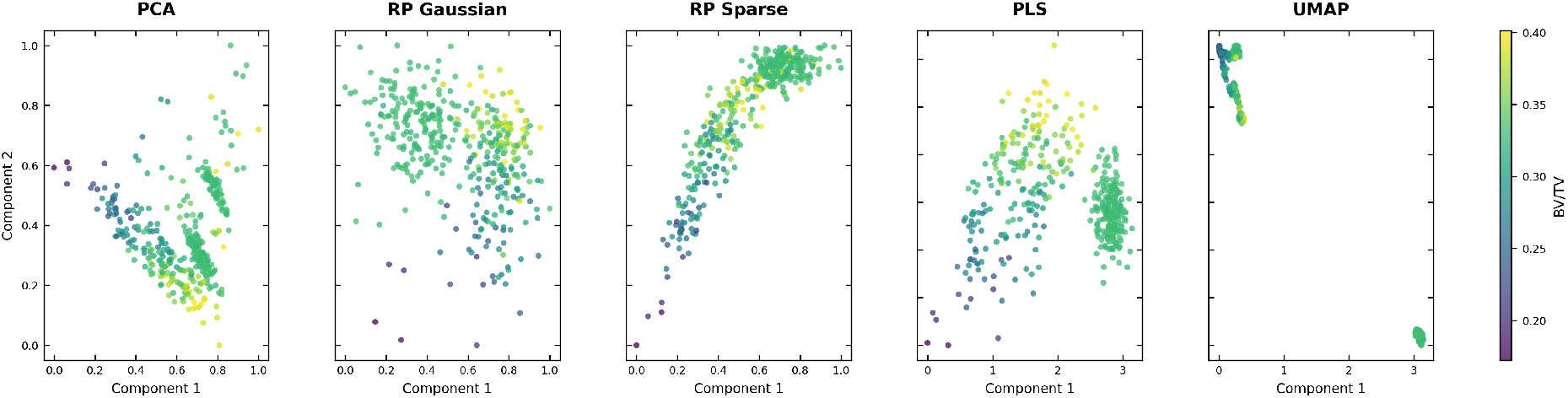
Two-dimensional projections of reduced feature spaces coloured by BV/TV. PCA, RP Gaussian, and RP Sparse produce diffuse point clouds with gradual colour gradients and substantial class overlap. PLS spreads points along a continuous supervised axis without sharp separation. UMAP (rightmost) forms tight, discrete clusters with clear BV/TV separation, consistent with its spectral clustering objective [10]. Only the first two components are shown for PCA, RP Gaussian, RP Sparse, and PLS (which use 8–16 components total); the 2D projection understates their information content.

This result contrasts with the repeated CV finding of +0.032 (quantum advantage, 19/25 folds). The sign reversal suggests that the positive trend observed under repeated CV was at least partially attributable to fold dependence: because repeated CV reuses data across folds, small favourable fluctuations for the quantum classifier on a single dataset can appear consistently across correlated fold estimates. The independent dataset design, which eliminates all such dependence, reveals that the UMAP quantum– classical gap is centred near zero and does not reliably favour either classifier.

Importantly, the UMAP gap under independent validation (−0.030) remains far smaller in magnitude than the deficits observed for linear methods under repeated CV (−0.090 to −0.116). While the independent validation was conducted only for UMAP, the linear method deficits were statistically significant even under corrected tests that account for fold dependence (Dietterich 5 × 2 CV *p* = 0.004 for PCA, *p* = 0.007 for PLS), making it unlikely that independent validation would eliminate those gaps. UMAP thus remains the only reduction method that preserves quantum kernel competitiveness, even though it does not confer a quantum advantage.

### 3.5 Tb.N Classification

Table 6 presents Tb.N classification results. Classical accuracies are near-ceiling (0.977–0.986), reflecting that Tb.N produces more easily separable texture signatures than BV/TV. Despite the compressed dynamic range, the relative pattern of quantum–classical gaps is preserved across methods, confirming the generalisability of the finding. UMAP again achieves the smallest quantum deficit (−0.001, statistically indistinguishable from zero), while all linear methods show gaps of −0.054 to −0.061. The consistency across both classification tasks — in which UMAP preserves quantum competitiveness while linear methods degrade it — constitutes the central finding of this work. Notably, the ordering among linear methods shifts between tasks (RP Gaussian is second-best for BV/TV but worst for Tb.N), reinforcing the importance of proper cross-validation and suggesting that differences among linear methods are not robust.

**Table 6:** Tb.N classification — 5 × 5 repeated stratified CV.

| Feature | Classical SVM | Quantum SVM (ZZ) | Gap |
| --- | --- | --- | --- |
| UMAP (8q) | $0.977 \pm 0.013$ | $0.976 \pm 0.013$ | $-0.001$ |
| PCA (8q) | $0.982 \pm 0.007$ | $0.928 \pm 0.015$ | $-0.054$ |
| PLS (4q) | $0.986 \pm 0.009$ | $0.930 \pm 0.020$ | $-0.056$ |
| RP Gauss. (8q) | $0.982 \pm 0.008$ | $0.921 \pm 0.021$ | $-0.061$ |

### 3.6 Quantum Kernel Ridge Regression

Table 7 presents quantum kernel ridge regression against classical Ridge baselines for continuous BV/TV prediction. All quantum regression *R*^2^ values are negative (worse than predicting the mean) except PLS (*R*^2^ = 0.185, using only 4 qubits). The ZZ feature map at 8 qubits maps 400 training points into a 2^8^ = 256-dimensional Hilbert space, producing a near-diagonal kernel matrix where most off-diagonal entries are close to zero. This effectively reduces to a lookup table rather than a smooth interpolator. PLS partially avoids this by using only 4 qubits (2^4^ = 16-dimensional Hilbert space), producing a denser kernel matrix, though still far below its classical counterpart (0.909). This classification–regression asymmetry suggests the ZZ quantum kernel captures categorical separability (enough for a decision boundary) but fails to encode the smooth distance relationships needed for metric regression.

**Table 7:**
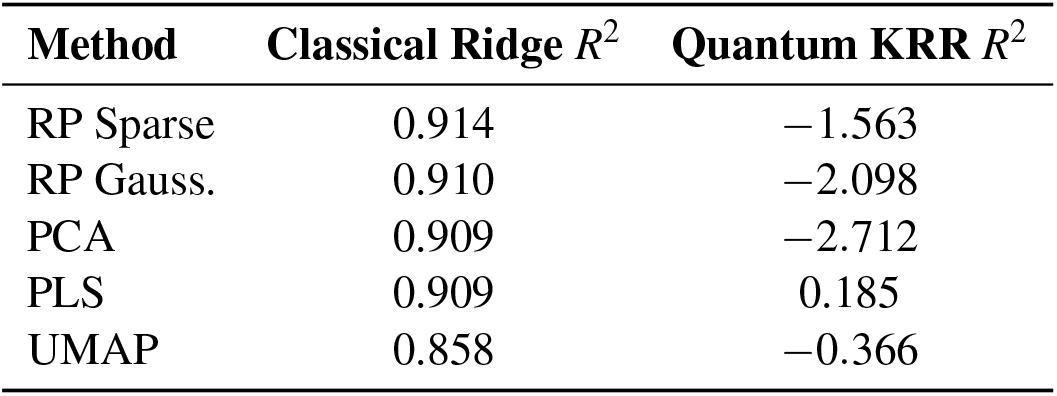
BV/TV regression — classical Ridge vs quantum KRR.

### 3.7 Kernel Matrix Diagnostics

Kernel matrix diagnostics (Table 8) provide quantitative confirmation. PCA and RP Gaussian produce near-diagonal quantum kernel matrices with effective ranks of 415.6 and 376.9 out of 500 (ratios 0.831 and 0.754) and off-diagonal means below 0.013. These kernels approach the identity matrix — a symptom of the exponential concentration phenomenon identified by Thanasilp et al. [4] — where each training point occupies a nearly orthogonal region of Hilbert space, leaving the SVM with no meaningful similarity structure to exploit. UMAP and PLS, by contrast, produce substantially denser kernels with effective ranks of 102.5 and 89.4 (ratios 0.205 and 0.179). UMAP’s kernel is particularly informative for classification: its off-diagonal mean (0.032) is modest, but its maximum off-diagonal entry (0.999) indicates near-identical kernel values within clusters and near-zero values between them — exactly the block-diagonal structure a binary SVM can exploit.

**Table 8:**
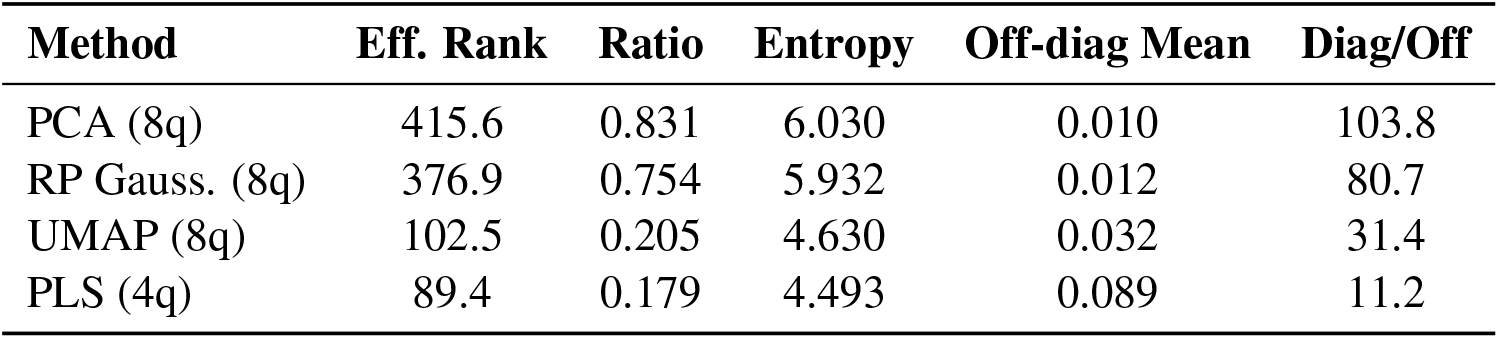
Quantum kernel matrix diagnostics.

## 4 Discussion

### 4.1 Synthetic Bone Generation

The zero-crossing Gaussian random field generator produced 500 synthetic trabecular bone volumes with 100% sample validity and perfect connectivity (LCC_raw_ = 1.000), confirming that the approach reliably produces plate-dominant architecture suitable for downstream analysis. Downstream Ridge regression *R*^2^ *≥* 0.87 across all reduction methods (Table 3) demonstrates that the 43 texture features extracted from synthetic slices encode substantial morphometric information, providing a meaningful test bed for comparing classical and quantum classifiers.

### 4.2 Nonlinear Reduction as Quantum-Compatible Preprocessing

The systematic comparison of five reduction methods reveals a clear nonlinear-vs-linear divide. UMAP — the only nonlinear, manifold-preserving reduction method tested — is also the only method that preserves quantum kernel competitiveness. All linear methods (PCA, RP Gaussian, PLS) produce substantial quantum deficits across both classification tasks, despite achieving higher downstream Ridge *R*^2^ (Table 3) and comparable or superior classical SVM accuracy. This dissociation between classical utility and quantum utility is key. Linear methods preserve global variance structure and pairwise distances, properties well-suited to classical RBF kernels. However, the ZZ feature map encodes inputs as rotation angles in a parameterised quantum circuit with entangling ZZ interaction gates, producing a quantum kernel that is periodic and local in its sensitivity: it responds strongly to small angular differences between nearby points but is relatively insensitive to global linear structure.

As Yang [10] proved, UMAP is equivalent to spectral clustering on the fuzzy *k*-nearest neighbourhood graph. This means UMAP embeddings are optimised to separate spectral clusters defined by the neighbourhood graph. The ZZ kernel’s periodic encoding can then distinguish between these pre-separated clusters without needing to discover the cluster structure itself. Kernel matrix diagnostics (Table 8) provide quantitative confirmation of this interpretation. This connects to the findings of Jiang and Otten [6], who showed that encoding strategy dominates quantum kernel performance by over 30% across datasets. Our results extend this principle to the preprocessing stage: even with a fixed encoding (ZZ feature map), the choice of reduction method produces accuracy variations of comparable magnitude (up to 0.105 between UMAP and PLS for BV/TV quantum classification). In the framework of Huang et al. [3], UMAP may succeed because it transforms the data geometry to better align with the quantum feature space.

### 4.3 Repeated CV versus Independent Validation

The independent dataset validation (Section 3.4) provides an important calibration of the repeated CV results. Under repeated CV, UMAP showed a +0.032 gap with the quantum kernel winning 19/25 folds and 9/10 folds under the 5 × 2 CV protocol. The independent validation, which eliminates all sources of fold dependence by using 10 separately generated datasets with independent reduction fits and kernel matrices, reversed the sign to −0.030 with the quantum kernel winning only 3/10 datasets (*t* = −1.70, *p* = 0.123; Wilcoxon *p* = 0.193; see Figure 3).

This discrepancy is consistent with the known optimistic bias of repeated cross-validation for corre-lated fold estimates. In repeated CV, training sets overlap substantially both within and across repeats; if the quantum classifier happens to benefit from a particular aspect of the single dataset’s geometry — for example, a fortuitous alignment between UMAP cluster boundaries and the ZZ kernel’s periodic structure — this advantage appears consistently across correlated folds, inflating the apparent win rate. The corrected tests (Nadeau–Bengio *p* = 0.168; Dietterich 5 × 2 CV *p* = 0.177) already indicated non-significance, but their correction factors are approximate. The independent validation provides a definitive answer: the UMAP quantum–classical gap is not reliably positive.

Nevertheless, the central finding of this work is preserved: UMAP keeps the quantum kernel competitive (gap near zero), whereas linear methods produce large, significant deficits. The independent validation refines the conclusion from “UMAP enables a possible quantum advantage” to “UMAP is the only reduction method that avoids a quantum penalty.” This is a practically important distinction: it identifies UMAP as a necessary preprocessing step for quantum kernel pipelines, even though it does not by itself confer a quantum advantage.

### 4.4 Classification vs Regression Failure

The uniform failure of quantum KRR (Table 7) reveals a fundamental limitation of the ZZ kernel at this scale and can be understood through the lens of exponential concentration [4]. Thanasilp et al. showed that quantum kernel values can concentrate exponentially towards a fixed value as the number of qubits grows, driven by at least four sources: embedding expressivity, entanglement, global measurements, and noise. In this regime, the estimated Gram matrix becomes uninformative — approaching the identity under the Loschmidt Echo test or a data-independent random matrix under the SWAP test — and the trained model’s predictions become independent of the input data. Although our statevector simulation eliminates the measurement noise pathway (we compute exact kernel values rather than statistical estimates), the underlying geometric concentration persists: the ZZ feature map with 2 repetitions and linear entanglement at 8 qubits produces a sufficiently expressive encoding that the fidelity between most pairs of data-encoded states becomes exponentially small. Our kernel matrix diagnostics (Table 8) provide direct empirical confirmation: PCA and RP Gaussian yield effective rank ratios of 0.831 and 0.754 with off-diagonal means below 0.013, indicating that the 500 × 500 kernel matrix closely approximates the identity.

This concentration manifests as a classification–regression asymmetry. For binary classification, the SVM requires only a separating hyperplane in the kernel-induced feature space — it suffices that same-class points share marginally higher kernel values than different-class points, even if all off-diagonal entries are close to zero. A near-identity Gram matrix with even weak block-diagonal structure can support a decision boundary. For regression, however, kernel ridge regression computes *α*_opt_ = (**K** + *λ* **I**)^−1^**y**, and when **K** *≈* **I** this reduces to *α*_opt_ *≈* **y***/*(1 + *λ*), producing a model that memorises training labels without learning any smooth relationship between inputs. Predictions on unseen test points then depend entirely on test-training kernel values, which are themselves near-zero, yielding predictions close to zero regardless of the true target value. This is precisely the data-independent prediction regime described by Thanasilp et al. (Corollary 1 in their framework), but arising here from geometric concentration of exact kernel values rather than from finite shot noise.

The PLS exception (4 qubits, *R*^2^ = 0.185) supports this interpretation. With only 4 qubits, the Hilbert space dimension is 2^4^ = 16, far below the 400 training points, which prevents the kernel from concentrating to the identity: the effective rank ratio drops to 0.179 and the off-diagonal mean rises to 0.089 (Table 8). This denser kernel retains enough inter-point similarity structure to enable partial regression, though it remains far below its classical counterpart (*R*^2^ = 0.909), likely because the periodic ZZ encoding still distorts the smooth metric relationships that linear Ridge regression exploits. Among the 8-qubit methods, UMAP achieves the least negative quantum *R*^2^ (−0.366 vs −1.563 to −2.712 for RP Sparse, RP Gaussian, and PCA), consistent with its lower effective rank ratio (0.205) and neighbourhood-preserving properties producing slightly more structured kernel matrices even when the overall kernel is near-diagonal.

These results have direct implications for QML pipeline design. At moderate qubit counts, quantum kernels can still classify — concentration degrades but does not eliminate the weak similarity contrasts needed for a decision boundary — but they cannot regress, because smooth interpolation requires meaningful off-diagonal kernel structure across the entire feature space. This makes quantum regression strictly harder than quantum classification at the same qubit count, not merely because the circuit is less expressive, but because the kernel matrix itself loses the metric structure that regression depends on. Practitioners should therefore verify kernel matrix density (effective rank ratio, off-diagonal mean) before attempting quantum regression, and consider reducing qubit count or using kernel designs less susceptible to concentration when continuous prediction is required.

### 4.5 Limitations and Future Work

Several limitations should be noted. All quantum kernels were computed via statevector simulation, which represents the theoretical best case — hardware noise is likely to exacerbate exponential concentration [4], pushing kernel matrices further toward the identity and potentially degrading classification accuracy. The evaluation uses synthetic rather than real micro-CT data, and only one nonlinear manifold method (UMAP) was tested. PLS results are additionally confounded by its lower qubit count (4 vs 8), and the fixed ZZ feature map may interact differently with reduction methods than variational encodings. The independent dataset validation was performed only for UMAP — future work should extend this to all reduction methods.

These constraints motivate several directions for future investigation. First, validation on real bone micro-CT data will determine whether the nonlinear-vs-linear divide observed here transfers to clinical imaging conditions. Second, evaluating other nonlinear reduction methods (e.g. t-SNE, autoencoders) will clarify whether UMAP’s advantage is specific to its spectral clustering properties or shared by nonlinear methods more broadly. Third, testing on quantum hardware with realistic noise models will reveal whether the quantum–classical parity observed for UMAP under ideal simulation survives device-level errors. Fourth, variational kernel optimisation may recover quantum competitiveness for linearly reduced features by adapting the encoding to the data geometry. Beyond static classification, the present framework lays the groundwork for a mechanics-aware quantum imaging approach: by extending these methods to time-series properties via quantum reservoir computing, we aim to process four-dimensional and higher-dimensional micro-CT data capturing mechanical deformation and damage evolution, ultimately toward the development of quantum image correlation strategies for materials characterisation.

## 5 Conclusion

We have shown that dimensionality reduction method choice critically determines whether quantum kernel SVMs remain competitive with classical baselines for trabecular bone classification. UMAP is the only method that avoids a quantum penalty, maintaining quantum–classical parity across two classification tasks and under independent dataset validation, while all linear methods produce substantial and significant quantum deficits. The ZZ quantum kernel fails uniformly at regression despite preserving competitiveness at classification, revealing a task-type asymmetry driven by near-diagonal kernel matrices. A novel zero-crossing Gaussian random field generator provides a controllable synthetic bone test bed with full morphometric validity. Together, these findings establish that manifold-preserving preprocessing is a necessary step for near-term quantum kernel pipelines and provide actionable guidelines for quantum feature engineering in biomedical imaging and materials characterisation.

## Acknowledgements

The authors acknowledge support from the School of Engineering and the School of Computing and Mathematical Sciences within the Faculty of Engineering and Science at the University of Greenwich. J.L.H. acknowledges support from Research England’s ‘Expanding Excellence in England’ grant via the “Multi-scale Multi-disciplinary Modelling for Impact” (M^3^4Impact) programme. E.A. acknowledges support from the National Quantum Computing Centre (NQCC) at the Rutherford Appleton Laboratory and the Yusuf Hamied Department of Chemistry at the University of Cambridge.

## Code and Data Availability

All code for synthetic bone generation, feature extraction, dimensionality reduction, quantum kernel computation, classical and quantum SVM classification, kernel ridge regression, independent dataset validation, and statistical analysis is publicly available at https://github.com/BellaZZ23/ quantum-bone-classification and archived on Zenodo [28] (DOI: 10.5281/zenodo.19831152). The synthetic datasets used in this study can be fully regenerated using the provided scripts with the random seeds specified in the code. The repository is released under the MIT licence.

## Notes

### Competing Interest Statement

The authors have declared no competing interest.

